# Unravelling termite evolution with 47 high-resolution genome assemblies

**DOI:** 10.1101/2025.01.20.633303

**Authors:** Cong Liu, Cédric Aumont, Alina A. Mikhailova, Tracy Audisio, Simon Hellemans, Yi Ming Weng, Shulin He, Crystal Clitheroe, Zongqing Wang, Ives Haifig, David Sillam-Dussès, Aleš Buček, Gaku Tokuda, Jan Šobotník, Mark C. Harrison, Dino P. McMahon, Thomas Bourguignon

**Author notes:** Corresponding authors: MCH; DPM; TB. These authors contributed equally: Cong Liu, Cédric Aumont, Alina Mikhailova, and Tracy Audisio. These authors contributed equally: Mark C. Harrison, Dino P. McMahon, and Thomas Bourguignon.

## Abstract

Termites are a lineage of social cockroaches abundant in tropical ecosystems where they are key decomposers of organic matter from wood to soil. Despite their ecological significance, only a handful of reference-quality termite genomes have been sequenced, which is insufficient to unravel the genetic mechanisms that have contributed to their ecological success. Here, we performed sequencing and hybrid assembly of 45 taxonomically and ecologically diverse termites and two cockroaches, resulting in haplotype-merged genome assemblies of 47 species, 22 of which were near-chromosome level. Next, we examined the link between termite dietary evolution and major genomic events. We found that Termitidae, which include ∼80% of described termite species, have larger genomes with more genes and a higher proportion of transposons than other termites. Our analyses identified a gene number expansion early in the evolution of Termitidae, including an expansion of the repertoire of CAZymes, the genes involved in lignocellulose degradation. Notably, this expansion of genomes and gene repertoires coincided with the origin of soil-feeding in Termitidae and remained unchanged in lineages that secondarily reverted to a wood-based diet. Overall, our sequencing effort multiplied the number of available termite genomes by six and provided unprecedented insights into the genome evolution of the most ancient lineage of social insects.

## Introduction

Termites are primarily known for their social lifestyle and wood-feeding habits. They live in colonies composed of a few hundred to several million individuals partitioned into several distinct castes: a working caste, comprising individuals that are sterile or temporarily give up reproduction; a generally sterile soldier caste; and a reproductive caste, represented by one or multiple kings and queens (Roisin, 2000; Roisin & Korb, 2011). Termites are one of the most abundant animal taxa in tropical and subtropical terrestrial ecosystems, where they are the primary decomposers of plant organic matter. They play a crucial role as ecosystem engineers (Eggleton et al., 1996; Evans et al., 2011; Jouquet et al., 2011; Ashton et al., 2019; Elizalde et al., 2020). Unlike ants, wasps, and bees (all Hymenoptera), termites are hemimetabolous diplo-diploid insects belonging to the order Blattodea, with the subsocial wood-feeding cockroaches of the genus *Cryptocercus* as the sister group (Lo et al., 2000). Termites first appeared in the fossil record in the Lower Cretaceous ∼130 million years ago (Mya) (Thorne et al., 2000; Engel et al., 2016). However, the bulk of termite diversity is found in the Termitidae, a family that originated during the Eocene around 50 Mya (Engel et al., 2011; Bucek et al., 2019), and which extensively diversified during the last 30 million years (Bourguignon et al., 2017; Wang et al., 2022). While 12 non-Termitidae families (or “lower” termites) feed only on sound wood or dry grasses with the aid of gut symbiotic protists, Termitidae include species lacking symbiotic protists that specialise in one type of organic matter quality along the wood-soil decomposition gradient (Donovan et al., 2001; Bourguignon et al., 2011). Because of their xylophagous behaviour, some species are also important pests of timber, accounting for a significant fraction of the total pest control costs in the regions where they occur (Gerozisis et al., 2008; Dhang, 2011; Rust & Su, 2012). Despite their ecological and economic importance in natural and urban areas, termites are poorly studied compared to Hymenopteran lineages of social insects.

So far, nine annotated termite genomes have been published. The first published termite genome was that of *Zootermopsis nevadensis* (Terrapon et al., 2014), a species selected for having a relatively small genome of approximately 560 Megabases (Mb), while other termites have genomes larger than 800 Mb (Koshikawa et al., 2008). The genome of *Z. nevadensis* was sequenced using short reads only and was highly fragmented, with a scaffold N50 of 740 kilobases (kb). Since then, short-read technology has been used to generate reference genomes of five additional termite species: *Macrotermes natalensis* (Poulsen et al., 2014), *Cryptotermes secundus* (Harrison et al., 2018), *Coptotermes formosanus* (Itakura et al., 2020), *Reticulitermes speratus* (Shigenobu et al., 2022), and *Reticulitermes lucifugus* (Martelossi et al., 2023). These genome assemblies were of slightly higher quality, with scaffold N50s varying between 1.18 and 2.00 Mb, except for the assembly of *R. lucifugus*, which had a scaffold N50 of 43 kb. It is important to note that these scaffolded short-read assemblies have many gaps, including up to 5000 Ns per 100 kb pairs (Harrison et al., 2018), with ungapped contig N50s ranging from 10.1 kb in *R. lucifugus* to 106.6 kb in *C. formosanus*. More recently, the genome of *C. secundus* was re-sequenced with PacBio High Fidelity reads together with the genomes of three new species: *Trinervitermes* sp. 1, *Odontotermes* sp. 2, and *Macrotermes bellicosus* (Qiu et al., 2023). As expected from long-read sequencing, the contiguity of these ungapped assemblies was substantially higher than previous termite genomes, with contig N50 ranging from 806 kb in *Trinervitermes* sp. 1 to 16.4 Mb in *M. bellicosus*.

The previously published termite genomes represent valuable resources. However, five of these nine genomes are extensively fragmented, while the most recent assemblies could benefit from additional scaffolding. This limitation in available high-quality resources hinders our understanding of termite genome evolution. The current standard for reference genome sequencing entails a hybrid genome assembly approach that combines long-read sequencing with Hi-C data (Dudchenko et al., 2017; Rhie et al., 2021). This has not been attempted for termites so far. In addition, the genomes of these nine species are not representative of termite taxonomic diversity, as only four of the 13 termite families and two of the 18 subfamilies of Termitidae (as currently recognised by Hellemans et al., 2024) are represented. Therefore, additional sequencing efforts are needed to generate a representative set of high-quality termite genomes. Here, we sequenced and annotated the genomes of 45 termite species alongside the cockroaches *Blatta orientalis* and *Cryptocercus meridianus*. Our sampling was representative of the taxonomic diversity of termites, including 11 of the 13 families of termites and 12 of the 18 subfamilies of Termitidae (Table S1), allowing us to identify the major events in genome evolution over the termite phylogenetic tree. Our sampling was also representative of the feeding diversity of Termitidae, including multiple lineages of wood feeders and multiple lineages of soil feeders, which account for a significant proportion of termite diversity but for which no genome reference was previously produced. This exhaustive sampling allowed us to determine how changes in diet have shaped termite genomes and influenced their diversity of carbohydrate-active enzymes (CAZymes) —the genes involved in the digestion of lignocellulose, the principal constituent of wood (Cantarel et al., 2009).

## Results and Discussion

### Expanding termite genome resources with 45 high-quality termite genomes

We sequenced 45 genomes of termite species belonging to 11 termite families and 12 subfamilies of Termitidae, as well as the genomes of two cockroach outgroups, *Blatta orientalis* and *Cryptocercus meridianus* (Table S1). For each species, we sequenced between 13.8 and 104.6 Gigabases (Gb) of PromethION long-read sequences (Table S2), which we used to generate the initial assemblies (Table S3). We also sequenced short reads for genome polishing, with a median genomic coverage of 32 (Table S4), and between one and nine transcriptomes per species (median = 3) for genome annotation (Table S5). Finally, we generated Omni-C and Hi-C data for the two cockroach species and 24 termite species. Omni-C and Hi-C data provide information on genome-wide chromatin interactions (Lieberman-Aiden et al., 2009), which we used to scaffold our genomes. Omni-C and Hi-C scaffolding produced near chromosome-level genome assemblies for 22 of the 26 species for which we produced these data. Our genome sequencing effort is the largest for termites so far, supplementing five termite genomes assembled with short reads only (Terrapon et al., 2014; Poulsen et al., 2014; Harrison et al., 2018; Itakura et al., 2020; Shigenobu et al., 2022; Martelossi et al., 2023) and four termite genomes assembled with PacBio High Fidelity reads only (Qiu et al., 2023) with an additional 45 genomes from diverse termite species.

The combination of sequencing approaches used in this study ensured the high quality of our 47 genomes (Figure 1). Before scaffolding, our long-read polished genome assemblies contained a median number of 3,981 contigs and had a median N50 value of 2.09 Mb, including 37 with an N50 greater than 1 Mb (Table S4). The median BUSCO completeness of polished genome assemblies was 99.2% based on the Insecta database (Table S4). In comparison, only three of the previously published termite genomes had a contig N50 greater than 1 Mb (Terrapon et al., 2014; Poulsen et al., 2014; Harrison et al., 2018; Itakura et al., 2020; Shigenobu et al., 2022; Martelossi et al., 2023; Qiu et al., 2023). Scaffolding with Omni-C or Hi-C markedly improved our genome assemblies: the 26 scaffolded genomes contained a median number of 2,253 scaffolds, had a median N50 of 44.1 Mb, and a median BUSCO completeness of 99.4% (Table S6). The quality of our genomes surpasses previously published termite genomes, and we significantly improved the genome assemblies of *Z. nevadensis* and *M. natalensis*, two previously sequenced termite species (Terrapon et al., 2014; Poulsen et al., 2014).

**Figure 1.**
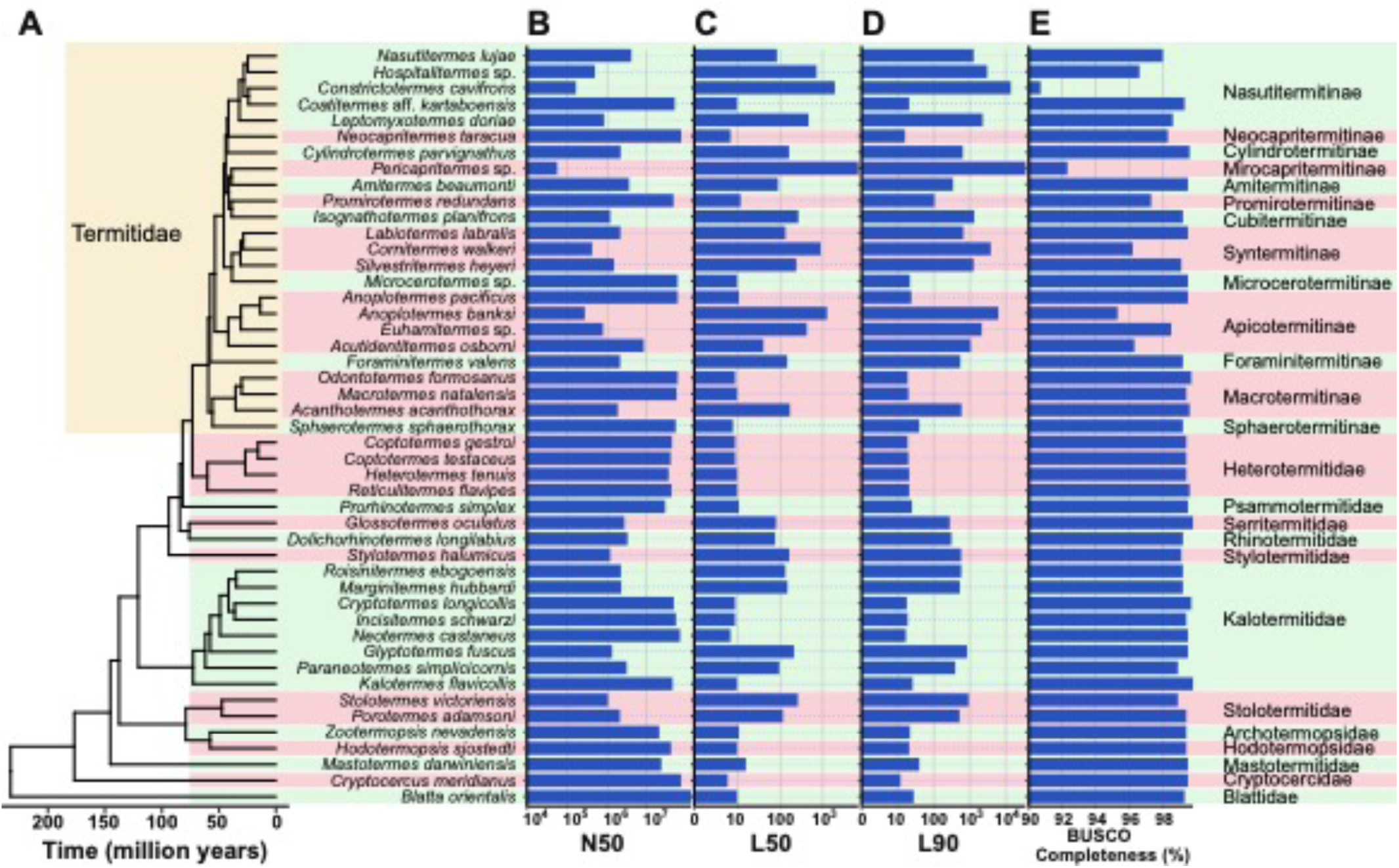
Assembly quality of the 47 genomes sequenced in this study. (A) Time-calibrated phylogenetic tree reconstructed with IQtree using 27,610 ultraconserved elements and calibrated with 1,409 single-copy orthologous genes, (B) N50, (C) L50, (D) L90, and (E) BUSCO Completeness.

### Termites and Cryptocercus share similar genomic features acquired before the origin of eusociality

The genome features of *Cryptocercus meridianus* were more similar to termites than to other cockroaches (Figure 2, Table S7). The size of the genome assembly of *C. meridianus* was 0.819 Gb, containing 15,909 gene models and 46.8% of transposons, falling within the range of values observed for termites (Genome size: 0.481-1.706 Gb; number of genes: 13,939-26,016; proportion of transposons: 24.2-48.5%). The only exception was the GC content of 38.2% in *C. meridianus*, which was lower than the 38.6% of GC found in *Zootermopsis nevadensis*, the termite species with the lowest GC content. In contrast, the genomes of other cockroaches were larger, contained more genes, and had a lower GC content. Our genome assembly of *Blatta orientalis* was 2.880 Gb, close to its estimated C-value of 2.963 Gb, and within the 2.083-5.046 Gb range of estimated C-values for non-*Cryptocercus* cockroaches (Koshikawa et al., 2008). The *B. orientalis* genome contained 55.8% of transposons, 31,064 gene models, and had a GC content of 35.5%, which was similar to the values previously reported for *Blattella germanica* and *Periplaneta americana*, having 55.8% and 62.9% of repeats, 28,575 and 29,939 predicted protein-coding genes, and 34.6% and 35.7% of GC content, respectively (Harrison et al., 2018; Li et al., 2018; Wang et al., 2022).

**Figure 2.**
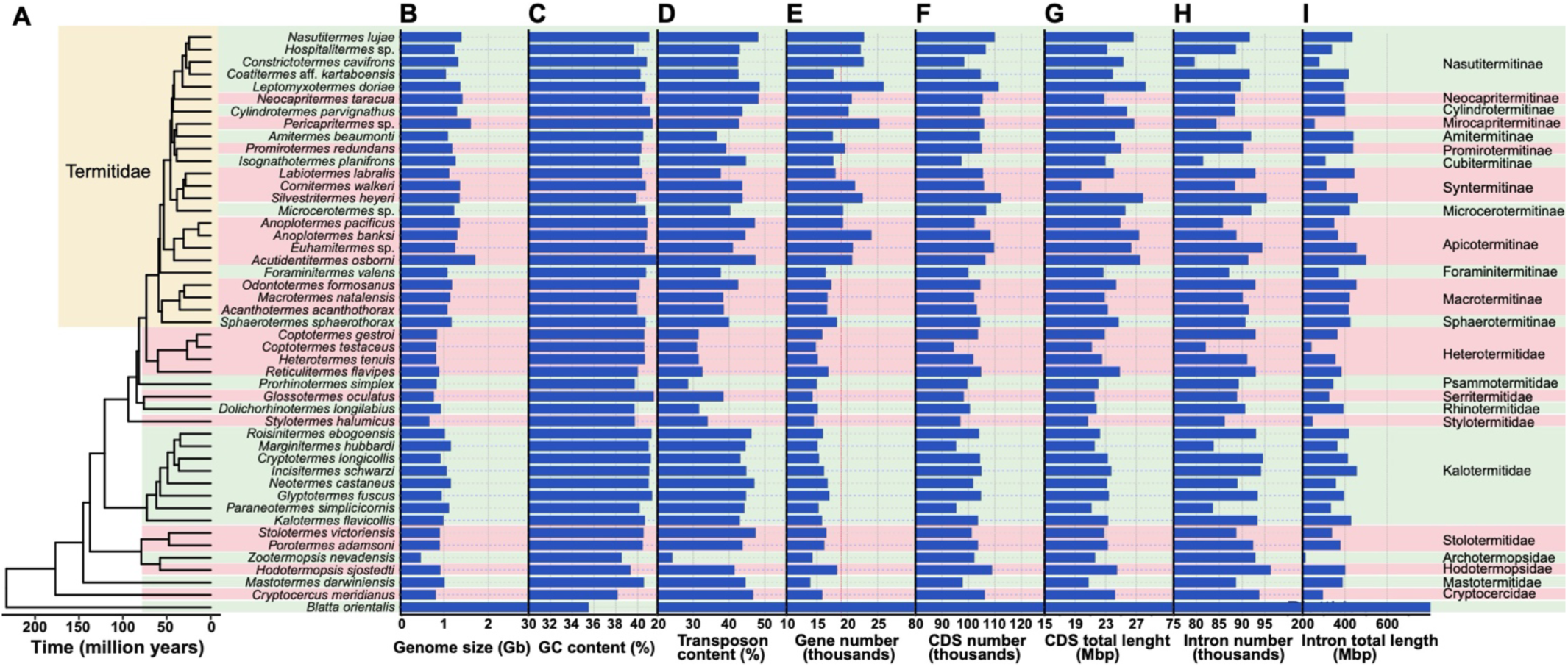
Characteristics of the 47 genomes sequenced in this study. (A) Time-calibrated phylogenetic tree reconstructed with IQtree using 27,610 ultraconserved elements and calibrated with 1,409 single-copy orthologous genes, (B) genome size, (C) GC content, (D) transposon content, (E) gene number, (F) CDS number, (G) total length of CDS, (H) intron number, and (I) total length. The red line corresponds to 19,000 genes.

We used evolutionary models to test whether the evolutionary dynamic of the genome of *Cryptocercus* resembles that of termites or cockroaches. We first reconstructed a robust maximum likelihood phylogenetic tree of *Blatta*, *Cryptocercus*, and termites using ultraconserved elements (UCEs) that we calibrated with 1,409 single-copy orthologous genes (Figures 1A, 2A). The topology and timing of our tree were largely consistent with previous phylogenetic trees based on thousands of nuclear markers (Bucek et al., 2019; Hellemans et al., 2024; Hellemans et al., 2022a), except for a few unresolved nodes differing in topology among analyses. We used this tree to model the evolution of eight genome characteristics: genome size, GC content, transposon content, gene number, number of coding sequences (CDS, which correspond to the number of exons without the untranslated regions), intron numbers, total coding CDS length, and intron total length. We constructed four lineage-dependent evolutionary models, each with two sets of parameters: two models separating *Blatta* + *Cryptocercus* from termites and two models separating *Blatta* from *Cryptocercus* + termites. The multi-rates models that considered *Blatta* + *Cryptocercus* in one regime systematically accrued <60% of the AIC model weight for each genome parameter, while the models using one set of parameters for *Cryptocercus* + termites accrued >95% of the AIC model weight for six genome parameters: genome size, gene number, CDS numbers, CDS total length, transposon content, and intron length (Figure 3A, Table S8). Therefore, AIC-weighted probabilities overwhelmingly supported the similarities of *Cryptocercus* and termite genomes. *Cryptocercus* and termites are sister groups and share multiple synapomorphies, such as biparental care of offspring, wood-feeding habits, or the symbiosis with gut lignocellulolytic protists helping them digest food (Cleveland et al., 1934; Nalepa & Jones, 1991). Our results showed that termites and *Cryptocercus* also have similar genomic features, including smaller genomes with fewer genes than other Blattodea, representing additional synapomorphies between the eusocial termites and the subsocial *Cryptocercus*. The similarities of *Cryptocercus* and termite genomes indicate that termites acquired their unique genomic features among Blattodea before they evolved eusociality.

**Figure 3.**
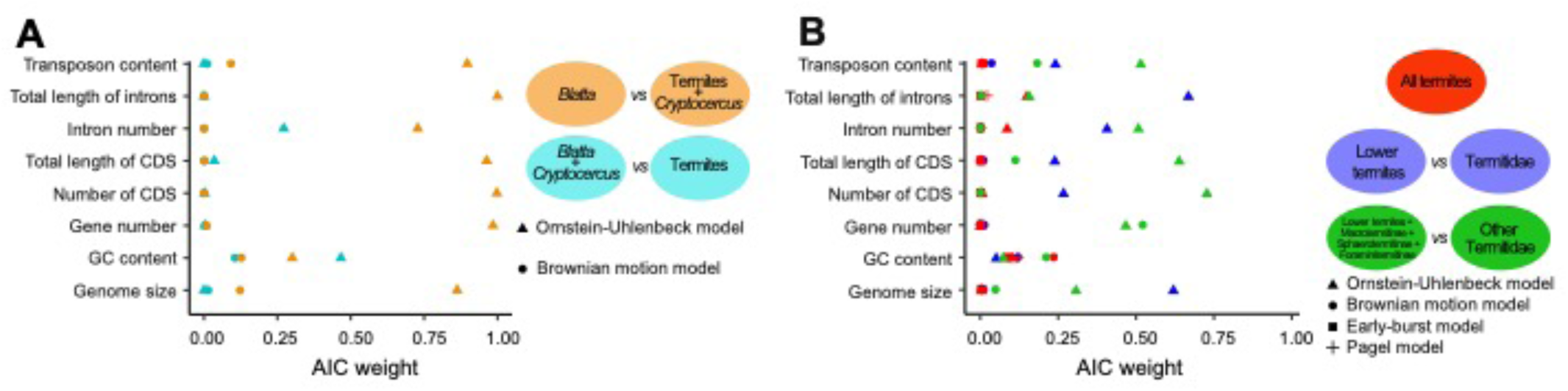
Evolution of genome characteristics in termites. (A) Total AIC weight of four multi-rate models reconstructing the evolution of eight genome characteristics in *Blatta*, *Cryptocercus*, and termites. Two models had distinct rates for *Blatta* and *Cryptocercus* + termites and followed a Brownian motion or an Ornstein-Uhlenbeck process. Two models had distinct rates for *Blatta* + *Cryptocercus* and termites and followed a Brownian motion or an Ornstein-Uhlenbeck process. (B) Total AIC weight of eight models reconstructing the evolution of eight genome characteristics in “lower” termites and Termitidae. Four models were multi-rate, with two distinct rates for “lower” termites and Termitidae or “lower” termites + Macrotermitinae + Sphaerotermitinae + Foraminitermitinae and other Termitidae, and followed a Brownian motion or an Ornstein-Uhlenbeck (OU) process. Four models were single-rate and followed a Brownian motion, an Early-burst model, an Ornstein-Uhlenbeck process, or a Pagel model.

### Termitidae genomes are the largest among termites and follow a unique evolutionary trajectory

The large number of termite genomes sequenced in this study allowed us to determine the main characteristics of their genomes (Figure 2, Table S7). The size of our 45 termite genome assemblies varied between 0.481 Gb and 1.706 Gb, which is in line with size estimations based on C-values that varied between 0.567 Gb and 1.858 Gb (Koshikawa et al. 2008). The overall transposon content varied between 24.2% and 48.5%, and the GC content showed slight variations, ranging from 38.6% to 41.7%. The genomes contained between 13,939 and 26,016 gene models, indicating that some species have larger proteomes than understood from the nine previously sequenced termite genomes, which contained a maximum of 18,162 gene models in *Cryptotermes secundus* (Harrison et al., 2018). Notably, the termite species with predicted proteomes exceeding 19,000 genes all belong to Termitidae and, more precisely, to the clade including all subfamilies of Termitidae except Macrotermitinae, Sphaerotermitinae, and Foraminitermitinae. These three termite subfamilies form two lineages branching before the rest of Termitidae, suggesting that the proteome size of the common ancestor of Termitidae was smaller than 19,000 genes. In summary, Termitidae appear to have larger genomes that generally contain more genes than other termites.

We tested whether the evolutionary dynamics of Termitidae genomes differ from other termite families, traditionally referred to as “lower” termites. We compared four lineage-independent mono-rate evolutionary models, in which the genome evolution of all termites is modelled with one set of parameters, with four lineage-dependent evolutionary models, in which termite genome evolution is modelled with two distinct sets of parameters for “lower” termites and Termitidae (two models) or “lower” termites + Macrotermitinae + Sphaerotermitinae + Foraminitermitinae and other Termitidae (two models). Lineage-independent models accrued <60% of the AIC model weight for all eight investigated genome parameter, while lineage-dependent models accrued >95% of the AIC model weight for five genome parameters: genome size, gene number, CDS number, CDS total length, and transposon content (Figure 3B, Table S9). Therefore, AIC-weighted probabilities indicated that these key genome traits differ significantly between “lower” termites and most Termitidae. Of note, the combined AIC model weight of the two models with distinct parameters for “lower” termites and Termitidae accrued >15% and the combined AIC model weight of the two models with distinct parameters for “lower” termites + Macrotermitinae + Sphaerotermitinae + Foraminitermitinae and other Termitidae also accrued >15% for all genome parameters except for gene number, for which the latter two models accrued a combined 98.8% of the AIC model weight. These results indicate uncertainties as to whether the genome evolution of Macrotermitinae, Sphaerotermitinae, and Foraminitermitinae evolved like those of the “lower” termites or other Termitidae for all key genome features but gene number, for which these families most certainly resemble “lower” termites. The changes in essential genomic features could be linked to the ecological success of Termitidae, the most diverse termite family. Notably, this family is also the only termite clade to entirely lack lignocellulolytic gut protists (Chouvenc et al., 2021).

### Termitidae have more genes and a higher proportion of transposons than other termites

We investigated how termite and *Cryptocercus* genome characteristics correlate with each other using our time-calibrated tree and Phylogenetic Generalised Least Square (PGLS). We focused on the eight genome characteristics displayed in Figure 2: genome size, GC content, transposon content, gene number, CDS number, CDS total length, intron number, and intron total length. Eighteen out of 28 PGLS analyses were significant with a False Discovery Rate (FDR) < 0.05 (Benjamini-Hochberg method), including five with a coefficient of determination (R-squared) higher than 0.5 (Figure 4, Table S10). Notably, nine of the ten non-significant correlations involved intron numbers and GC content, indicating that these two genomic features are largely independent of the other six. The strongest correlation was between genome size and transposon content (R-squared = 0.909; PGLS p-value < 10^-6^), indicating that the primary factor responsible for genome size variation in termites is transposons, as previously described for many organisms (Canapa et al., 2016; Elliott et al., 2015). The second strongest correlation was between genome size and gene number (R-squared = 0.650; PGLS p-value < 10^-6^), which, however, contrasts with the weak correlation between genome size and CDS number (R-squared = 0.156; PGLS p-value = 0.007) and the lack of correlation between genome size and intron number (R-squared = 0.012; PGLS p-value = 0.160). These results indicate that more gene models were predicted for termites with larger genomes while the number of CDS (or exons) was less strongly affected. In other words, more gene models containing fewer exons were predicted for termites with larger genomes (Figure 5A-D). Current genome annotation methods are imperfect and prone to errors, as illustrated by the human genome, for which the exact number of genes is still unknown (Guigó, 2023). Some genomic features, such as repetitive elements, affect the number of predicted gene models (Tørresen et al., 2019). We reasoned that the larger and more repetitive (Termitidae) genomes may be more affected by this type of annotation error than comparatively smaller genomes, potentially leading to genome-size-dependent bias in the number of false gene predictions. To verify the positive correlation between genome size and gene number and minimise the potential influence of an annotation bias, we performed additional PGLS analyses between genome size and four other estimates of gene number: the total number of genes belonging to hierarchical orthologous groups (HOGs) and the total number of genes longer than 100, 150, and 250 amino acids. The reasoning is that gene models not belonging to HOGs and shorter gene models might be more abundant in larger termite genomes due to annotation bias. However, all four PGLS analyses remained significant with R-squared values > 0.5 (Figure 5E-H, Table S11), validating the correlation between genome size and gene number. Therefore, termites with larger genomes, particularly those within the clade including all subfamilies of Termitidae except Macrotermitinae, Sphaerotermitinae, and Foraminitermitinae, which are the most ecologically diverse and species-rich clade of termites, appear to have a higher number of genes than other termites.

**Figure 4.**
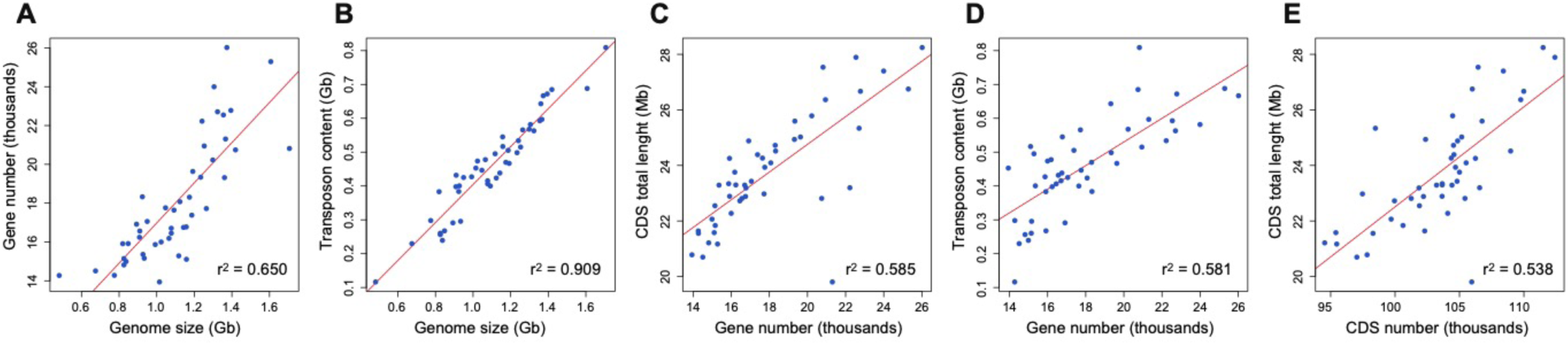
Relationships among the main genome characteristics of termites and *Cryptocercus* having a coefficient of determination (R-squared) higher than 0.5. Phylogenetic Generalised Least Square (PGLS) regressions between (A) genome size and gene number, (B) genome size and transposon content, (C) gene number and total length of CDS, (D) gene number and transposon content, and (E) CDS number and CDS total length.

**Figure 5.**
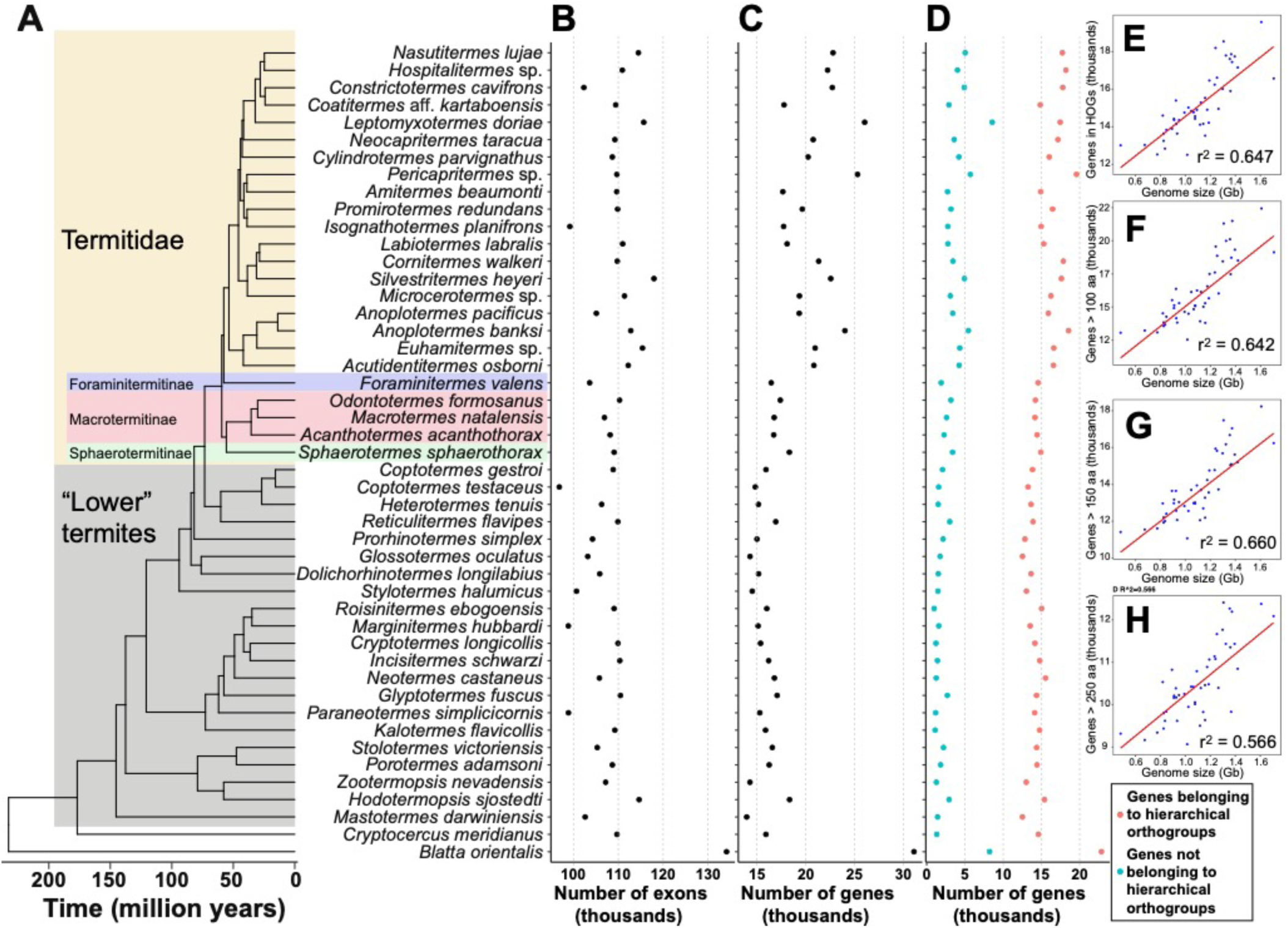
Number of predicted gene models and their relationship with genome size. (A) Time-calibrated phylogenetic tree reconstructed with IQtree using 27,610 ultraconserved elements and calibrated with 1,409 single-copy orthologous genes, (B) number of exons, (C) number of genes, (D) number of genes belonging or not to hierarchical orthogroups (HOGs), and relationship between genome size and (E) number of genes belonging to HOGs, (F) number of genes longer than 100 amino acids, (G) 150 amino acids, and (H) 250 amino acids.

### Diet does not affect termite genome features

Modern termite families have retained the wood diet of their common ancestor, except for many Termitidae that have a soil-feeding ancestor and feed on soil or secondarily reacquired a wood diet (Bourguignon et al., 2011; Donovan et al., 2001; Hellemans et al., 2022b). Our sampling included wood feeders from 11 of the 13 termite families and representatives of multiple lineages of soil feeders within the Termitidae, allowing us to investigate the effect of diet on the evolution of termite genomes. We first used a phylogenetic ANOVA to compare wood-feeding termites + *Cryptocercus* to soil-feeding termites for eight genomic features. Four genomic features, GC content, CDS number, intron number, and intron total length, did not significantly differ between wood and soil feeders, while the other four genomic features, genome size, transposon content, gene number, and total length of CDS, showed significantly higher values in soil feeders than wood feeders (Tables S7, S12). However, because all soil-feeding termites belong to Termitidae, these initial results may reflect the unique evolutionary trajectory of Termitidae among termites rather than the influence of diet on genome evolution. In support of this hypothesis, no comparisons were significant in phylogenetic ANOVAs comparing wood-feeding Termitidae only to soil-feeding termites (Table S12). Therefore, while the ancestor of non-Macrotermitinae non-Sphaerotermitinae non-Foraminitermitinae Termitidae experienced an expansion of genome size and gene number around the time they switched from a wood-based diet to a soil-based diet, diet had no apparent effect on the main genomic features in Termitidae species that switched back to wood-feeding.

### Termites have a reduced CAZyme repertoire compared to cockroaches

Although the ability of termites to decompose wood largely depends on their association with symbiotic gut microbes (Brune, 2014), they also produce their own cellulases (Tokuda et al., 2004; Watanabe et al., 1998). We searched our 45 termite and two cockroach genomes for Carbohydrate Active Enzymes (CAZymes) belonging to glycosyl hydrolases (GHs), carbohydrate esterases (CEs), and auxiliary activities (AAs), which are involved in the degradation of lignocellulose (Cantarel et al., 2009), a recalcitrant polymer composed of lignin, cellulose, and hemicellulose (Lynd et al., 2002). We also searched for their associated carbohydrate-binding modules (CBMs) when present, which bind to carbohydrates and glycoconjugates, but we did not search for glycosyl transferases (GTs), as they participate in the formation of glycosidic bonds rather than their degradation (Coutinho et al., 2003). Each termite genome contained between 151 and 213 CAZymes involved in the degradation of carbohydrates and glycoconjugates from 44 families, including 30 GHs, three CEs, three AAs, and eight CBMs (Figure 6A-B, Table S13). Out of these 44 CAZyme families, 28 were present in all termite genomes and eight in the genomes of all but one species (Table S13), indicating that the CAZymes previously found in the genomes of a few species (He et al., 2023; Zhang et al., 2012) are universally part of the genomic CAZyme repertoire of termites. Some CAZymes encoded in termite genomes are not involved in lignocellulose degradation, such as GH18, which targets chitin and was represented by 12 to 29 copies per genome, and CBM14, which specifically binds to chitin and was represented by 14 to 24 copies per genome. These CAZymes are likely involved in the biosynthesis and recycling of the peritrophic matrix lining the midgut of termites (Fujita et al., 2010; Morales-Ramos et al., 2006; Sandoval-Mojica & Scharf, 2016), the development and degradation of the chitin matrix in epithelia during moulting (Rabadiya and Behr, 2024), or may be part of the host defence mechanism against fungi and other chitin-containing pathogens.

**Figure 6.**
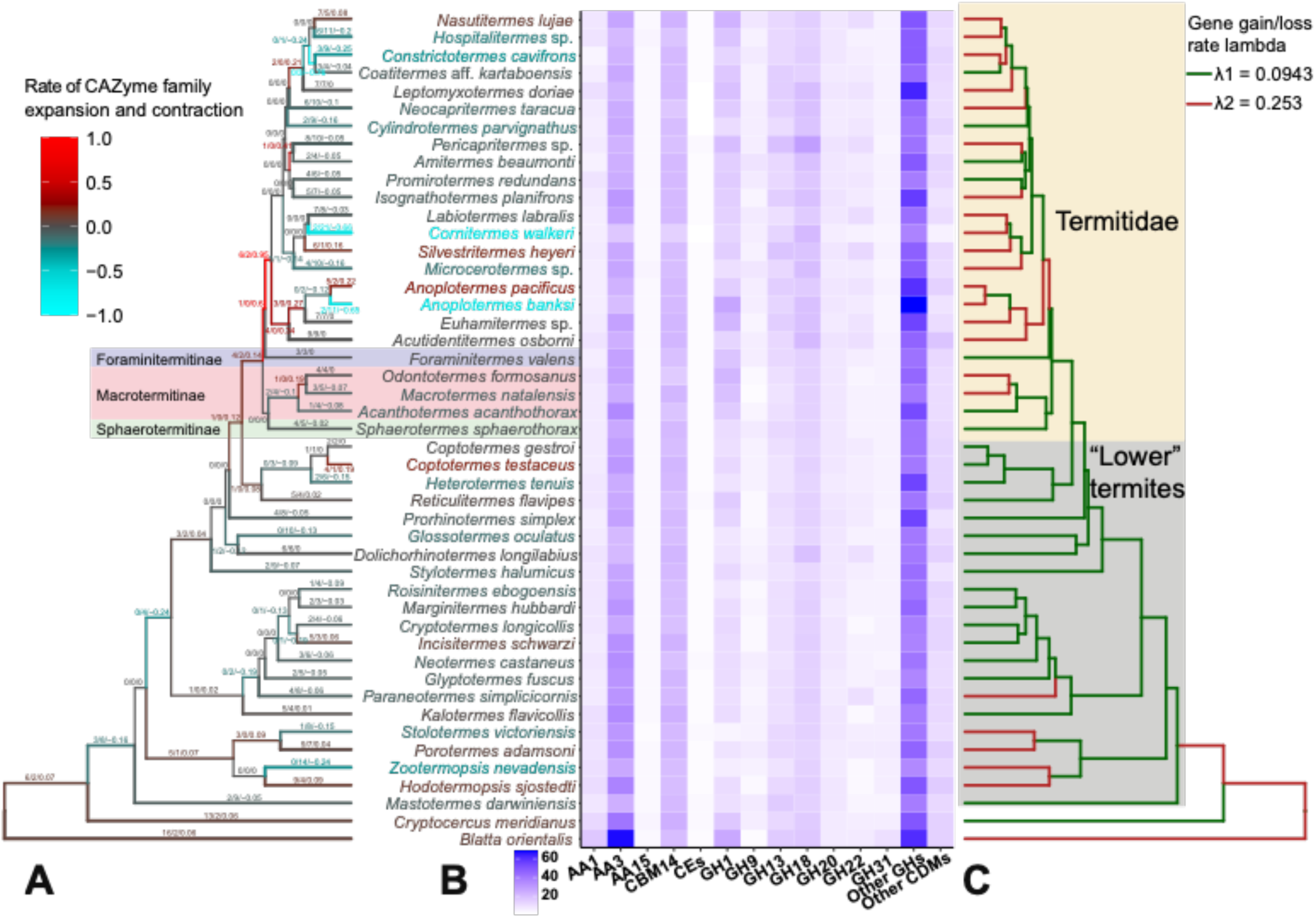
Evolution of the CAZyme repertoire of the 47 genomes sequenced in this study. (A) Time-calibrated phylogenetic tree showing the number of CAZyme families found to significantly expand (red) and contract (blue) with CAFE. The three successive numbers of the branch labels are the number of gene families significantly expanding, the number of gene families significantly contracting, and the overall expansion (or contraction) rate per million years. (B) Heat map showing the abundance of the CAZyme families across genomes. (C) Time-calibrated phylogenetic tree showing branches with low (green) and high (red) average gene gain/loss rates, Lambda, across all gene families.

We searched for CAZymes in the genomes of *C. meridianus* and *B. orientalis* and found that they respectively contained 218 and 269 CAZymes involved in the degradation of carbohydrates and glycoconjugates. The families dominating the CAZyme repertoire of cockroaches were largely those also found in the repertoire of termites (Figure 6B, Table S13). For example, we found 20 copies of CBM14 and 17 of GH18 in *B. orientalis* and 24 copies of CBM14 and 16 of GH18 in *C. meridianus*. Therefore, the CAZyme repertoire of cockroaches and termites is largely conserved in terms of family representation but not in terms of number of CAZyme genes. This is well-illustrated by the largest CAZyme repertoire among all analysed termite genomes, the CAZyme repertoires of *Anoplotermes banksi* (213 CAZymes), which is itself smaller than either sequenced cockroach. The reduced CAZyme repertoire of termite genomes compared to cockroach genomes may stem from several factors, including the development of social immunity and the reliance of termites on their gut microbes to digest wood at various stages of decomposition.

### The genomic CAZyme repertoire of Termitidae expanded, especially in soil-feeding species

The size of the CAZyme repertoire varied between 151 and 213 CAZymes per termite genome. We investigated the dynamic of this size variation across termites using CAFE5 (Mendes et al., 2020), which reconstructed the history of CAZyme gene family expansions and contractions along our termite time-calibrated phylogenetic tree. We focused on the 39 CAZyme families present in at least 20 genomes. The model with the highest likelihood was a model with two gamma-distributed rate categories *K* assigned to gene families and two local gene loss and gain lambda parameters assigned to branches of the phylogenetic tree. Among the 39 CAZyme families, 18 were assigned a gamma rate of 1.696 and considered fast evolving, 20 were assigned a gamma rate of 0.304 and considered slow evolving, and one did not fall into either of the two categories (Table S14). Notably, the average lambda, which indicates the rate of gene loss and gain along each branch of the tree, was higher (λ2 = 0.253) in many branches of Termitidae and several branches outside the Kalotermitidae + Neoisoptera than in the rest of the tree (λ1 = 0.0943) (Figure 6C). In addition, our analyses identified an accelerated expansion of the CAZyme repertoire early in the evolution of Termitidae, especially at the origin of the clade including all subfamilies of Termitidae except Macrotermitinae, Sphaerotermitinae, and Foraminitermitinae (Figure 6A). The expansion of the CAZyme repertoire in Termitidae, and more specifically in Termitidae excluding Macrotermitinae, Sphaerotermitinae, and Foraminitermitinae, coincides with the overall expansion of the number of genes and changes in genomic features and genome evolutionary dynamics observed for this node. Termitidae comprise about three-quarters of described termite species (Krishna et al., 2013) and are the most ecologically diverse termite family, including species feeding along the entire plant organic decomposition gradient, from wood to soil (Bourguignon et al., 2011; Donovan et al., 2001; Hellemans et al., 2022b). The ecological success of Termitidae was accompanied by an expansion of the total number of genes, including an expansion of the CAZyme repertoire.

While diet did not affect the main genomic features in modern termites, it may have affected the CAZyme repertoire. We compared the size of the CAZyme repertoire of *Cryptocercus* and termite species feeding on wood to that of termite species feeding on soil using a phylogenetic ANOVA and found that soil feeders encoded more CAZyme genes in their genomes than wood feeders (p = 0.009, Table S15). The difference remained significant in the phylogenetic ANOVA performed on species belonging to Termitidae only (p = 0.011, Table S15). However, phylogenetic ANOVA performed on each CAZyme family separately did not allow us to identify which CAZyme families are more abundant in soil feeders than in wood feeders (Table S15). Soil feeders primarily utilise the protein-rich fraction of the soil, including soil microbes and the organic residues that bind to clay particles (Ji & Brune, 2001, 2005), while lignocellulose is scarce in their diet (Brune, 2014). The cause of the expansion of the CAZyme repertoire in soil feeders is thus unclear from the perspective of diet processing and may be linked to other factors, such as the inactivation of the elevated pathogen loads in the diet and environment of soil feeders.

### Conclusion

Termites have historically received less attention from researchers than social insects belonging to Hymenoptera, which include bees, ants, and wasps. Their genomes have also been comparatively less studied. In this study, we sequenced, annotated, and compared the genome characteristics of 45 termite species representing the ecological and taxonomic diversity of termites, including 11 of the 13 extant families and 12 of the 18 extant subfamilies of Termitidae. We also sequenced two cockroach outgroups, *Blatta orientalis* and *Cryptocercus meridianus*. The overall similarity of termite and *Cryptocercus* genomes, which are smaller and encode fewer genes than other cockroach genomes, indicate that termites acquired their unique genome features before the origin of eusociality, raising thought-provoking questions about the nature of the evolutionary changes that underpinned termite social transitions. Within termites, the analysis of our 45 genomes highlights the peculiarity of Termitidae, the most species-rich and ecologically diverse termite family, which experienced an expansion in the size and gene content of their genomes. Overall, our results shed new light on the evolution of termites from a largely unexplored angle: their genomes.

## MATERIALS AND METHODS

### Sampling

We used samples from 45 species of termites and two species of cockroaches that are representative of the global termite diversity (Table S1). Samples of every species were collected in the following preservative agents: 80% ethanol for morphological species identification; RNA-later®, 100% ethanol, or live specimens for DNA extraction; TRIzol^TM^ reagent or live specimens for RNA extraction; and live specimens for Omni-C and Hi-C library preparation. For each termite species sequenced in this study, we only collected individuals coming from a single colony, with a few exceptions for samples collected for RNA sequencing required for genome annotation (Table S5). We preferentially used specimens from termite colonies maintained in lab breeds, as it ensured continuous access to high-quality material. The two cockroach genomes were generated using a single individual preserved in RNA-later® and originating from laboratory breeds for *C. meridianus* and *B. orientalis*, respectively. For species not kept in laboratory, samples were collected in the field and stored in RNA-later®, 100% ethanol, and TRIzol^TM^ reagent at -80°C until DNA and RNA extraction.

### DNA extraction

We extracted high molecular weight DNA from 45 species of termites and two species of cockroaches either using the Qiagen DNeasy® Blood and Tissue kit, the MagAttract HMW DNA kit, or the Nanobind Tissue kit from PacBio®. For most species, in order to obtain sufficient DNA yield for the preparation of both long- and short-read DNA libraries, we pooled the heads of one to twelve worker specimens sampled in the same colony. We used one king and one queen head for *Macrotermes natalensis*. For four species, we extracted DNA from different tissues: from the head, muscles, and legs for *C. meridianus*; from muscles of a single queen for *Anoplotermes pacificus*; from one nymph head and one nymph without gut for *Reticulitermes flavipes*; and from the head and all legs of a worker for *Mastotermes darwiniensis*. We used dissected worker heads and termite tissues without guts to avoid any potential contamination from gut microbes. The same DNA extracts were used for the short-read and long-read DNA library preparation to avoid any potential downstream assembly issues.

### Whole-genome sequencing with long reads

The long-read DNA libraries were prepared and sequenced by the DNA Sequencing Section of the Okinawa Institute of Science and Technology (OIST), Japan, and by the DRESDEN-concept Genome Center (DcGC), Germany. Quality checks were performed prior to sequencing with a Qubit Flex Fluorometer and a NanoDrop 2000C to ensure high-quality DNA extracts. The size distribution of nucleic acids was determined with an Agilent Femto Pulse. Samples with high size distributions (>50 kb) were mechanically sheared with a Megaruptor® 3 and sequenced with the Oxford Nanopore PromethION platform. Between 10 and 50 fmol of libraries were loaded on an Oxford Nanopore PromethION flowcell, and the specified run length was 72 h.

### Whole-genome sequencing with short-reads

The remaining DNA extracts not used for long-read sequencing were used to produce short-read sequence data for genome polishing. For samples processed at OIST, library preparation was performed using the NEBNext Ultra II FS DNA Library Prep Kits with one-fifth of the recommended reagent volumes. Paired-end 150 bp sequencing was performed on the Illumina HiSeq X platform. For samples processed at DcGC, linked-read libraries were prepared for nine species with the Tell-Seq WGS Library Prep Kit following the manufacturer protocol. Paired-end 150bp sequencing of linked-read libraries was performed using the NovaSeq 6000 platform with a S4 XP v1.5 Flowcell. Note that the short reads prepared with TELL-Seq were only used for polishing.

### Omni-C and Hi-C library preparations and sequencing

We performed Omni-C and Hi-C library preparations for 24 termite species and two cockroach species. For each termite species, chromatin isolation was performed using one to ten heads dissected from one live worker, except for *Anoplotermes pacificus* and *Macrotermes natalensis*, for which legs and muscles from other body parts were used. For the two cockroach species, chromatin was isolated from dissected heads or legs. Dissected tissues were snap-frozen and deposited in a foil-lined mortar pre-cooled with liquid nitrogen. Head tissues were ground into a fine powder using a pestle and carefully transferred from the foil into a pre-cooled tube for chromatin digestion. At OIST, the Omni-C library preparation was performed following the standard Dovetail® Omni-C® protocol. At DcGC, the Hi-C library preparation was performed using the KAPA HyperPrep Kit and the Arima-HiC 2.0 Kit following the manufacturer’s protocols. Paired-end 150 bp sequencing was performed on the Illumina HiSeq X platform or the NovaSeq 6000 platform.

### Transcriptome sequencing

We generated a series of transcriptomes derived from various life stages (egg, larva, and adult imago), castes (worker, soldier, and primary reproductive), and body parts (head, whole body with or without gut, and gut) for all genomes sequenced here (Table S5), except for the genome of *Zootermopsis nevadensis*, for which we used 26 transcriptomes. We attempted to sequence as many transcriptomes as allowed by material availability for every species. All transcriptomes were generated to annotate the genomes. Total RNA was extracted using the Invitrogen^TM^ PureLink^TM^ RNA Mini kit at OIST and the RNeasy® Plus mini kit (Qiagen, Germany) at DcGC. Messenger RNA (mRNA) enrichment was performed using the NEBNext® Poly(A) mRNA Magnetic Isolation Module. Reverse transcription of mRNA into cDNA was performed with the NEB Ultra II Directional RNA Library Prep Kit. Libraries were prepared using the NEBnext® Ultra™ II RNA Library Preparation kit for Illumina®. Paired-end 150 bp sequencing was performed on the Illumina HiSeq X or the NovaSeq 6000.

### Genome assembly

We used Nanopore long-read data to generate the initial genome assemblies. General statistics about raw long-read data were generated using NanoStat v1.0 (De Coster et al., 2018) to assess read quality. The long-read statistics of every species sequenced here are available in Table S2. Long reads encompassing protein-coding regions were frameshift-corrected by mapping proteins of the Uniprot database (accessed on June 16, 2021) to long reads using Proovframe v0.9.7 (Hackl et al., 2021) and DIAMOND v2.0.4.142 (Buchfink et al., 2015) both used under default settings (Table S2). Long reads were assembled into contigs using Flye v2.8.1 and v2.9 (Kolmogorov et al., 2019) with default settings.

We used Illumina short-read data to polish the genome assemblies obtained with long reads, an essential step given the high error rates of long-read data (see Table S2). For species sequenced at DcGC, TELL-seq reads were used and processed with Tell-Read v1.0.3 (Chen et al., 2020). Short paired-end reads were first checked for contaminations. Briefly, short reads were trimmed from adapters, low-quality bases, and trailing polyG tails using fastp v0.20.1 under default settings (Chen et al., 2018). Trimmed reads were downsampled using the BBNorm tool from BBMap v38.86 (Bushnell, 2014) to 50x coverage based on kmer size of 31. Short reads were downsampled to avoid misassemblies. Short reads were assembled with metaSPAdes v3.15.1 (Nurk et al., 2017) settings with kmer sizes of 21, 31, 41, 51, 71, and 91. Mitochondrial genome fragments were identified and annotated using MitoFinder v1.4 (Allio et al., 2020). Each fragment was searched against an in-house termite mitochondrial genome database and a database composed of all mitochondrial genome fragments from the 47 species used in this study with megaBLASTn from the BLAST+ v2.13.0 suite of software (Camacho et al., 2009). Libraries containing foreign mitochondrial contigs (contaminants) were discarded. This procedure ensured the absence of contaminations in the short reads used in this study. Following the confirmation of the absence of contamination, short reads were mapped to long-read genome assemblies using minimap2 (Li, 2018), with all parameters left as default. The resulting binary alignment maps were used to polish the draft assemblies using a k-mer-based binning approach with HyPo v1.0.3 (Kundu et al., 2019), run with default parameters. Low-coverage regions were removed, and the assemblies were collapsed into a single haplotype using three successive commands of purge_haplotigs v1.1.2 (Roach et al., 2018). First, a coverage histogram was generated with the command “purge_haplotigs hist”. Second, the coverage was analysed on a contig-by-contig basis using the command “purge_haplotigs cov -l X -m Y -h Z -j 80 -s 80”, where the values of X, Y, and Z varied among genomes. Third, the purging pipeline was run using the command “purge_haplotigs purge -a 80”. Other parameters of the purge_haplotigs command not mentioned above were left as default.

We scaffolded the polished assemblies of 26 species using Omni-C and Hi-C read data. The Omni-C and Hi-C reads were mapped onto each corresponding genome using BWA-MEM (Li, 2013) under default parameters. Following this mapping step, paired-end reads of Omni-C and Hi-C were processed using pairtools v1.0 (Open2C et al., 2024). Briefly, Omni-C and Hi-C reads were arranged using the block-sorting approach with the command “pairtools sort --nproc 16” and PCR and optical duplicates were removed with the command “pairtools dedup --nproc-in 8 --nproc-out 8 --mark-dups”. Mapped read pairs were used to generate scaffolds using YAHs v1.1 (Zhou et al., 2023) with all parameters left as default for scaffolding with Omni-C and Hi-C data and with the command “-e GATC,GANTC,CTNAG,TTAA” specifying the restriction enzymes for the scaffolding with Hi-C data. Assembly gaps present in the scaffolds were filled using long reads with one iteration of LR-Gapcloser v1.0 (Xu et al., 2019) set on nanopore reads and other parameters left as default.

### Genome assembly evaluation

The draft assemblies were evaluated at each assembly step by assessing the total length of the assembly, contig number, contig length, and N50 and L50 values with QUAST v5.0.2 (Gurevich et al., 2013) (see Tables S3-4, S6). Genome completeness was estimated at each assembly step by counting the number and completeness of 1367 core insect genes with BUSCO v5.0 (Seppey et al., 2019), using the command line “busco --lineage_dataset insecta_odb10 --mode genome”. The quality of the genome assemblies was assessed with MERQURY v1.3 (Rhie et al., 2020).

### Genome annotation

We annotated the 45 genomes of termites and two genomes of cockroaches sequenced in this study using an in-house pipeline combining the outputs of several annotating software. All software was run using default settings. For each genome assembly, repeat elements were detected using RepeatModeler v.2.0.3 (Smit et al., 2015b) and masked with RepeatMasker v.4.1.2 (Smit et al., 2015a). Protein-coding genes were identified in the masked genome assemblies using a combination of protein-to-genome alignments, transcript-to-genome alignments, and *ab initio* gene predictions.

First, we obtained extrinsic evidence for the annotation of protein-coding genes using both protein-to-genome and transcript-to-genome alignments. (1) We used the proteomes of ten model insect species (*Aedes aegypti*, *Anopheles gambiae*, *Drosophila melanogaster*, *Acyrthosiphon pisum*, *Apis mellifera*, *Nasonia vitripennis*, *Solenopsis invicta*, *Bombyx mori*, *Manduca sexta*, and *Tribolium castaneum*), three termites (*Cryptotermes secundus*, *Zootermopsis nevadensis*, and *Coptotermes formosanus*), and one cockroach (*Blattella germanica*) for protein-to-genome alignments. The proteomes of the ten model insects were retrieved from NCBI. The proteomes of the three termites and the cockroach were retrieved from InsectBase (Mei et al., 2022). Protein sequences were aligned to every 47 masked genomes with miniprot v.0.7 (Li, 2023). (2) We used the transcriptomes sequenced together with the 47 genomes to generate transcript-to-genome alignments (Table S5). RNA reads were mapped onto each masked genome with Hisat2 v.2.2.1 (Kim et al., 2019). Mapped RNA reads were used to generate genome-guided transcriptome assembly with StringTie v.2.1.4 (Pertea et al., 2015).

Second, we used three *ab initio* gene predictors. The first predictor, AUGUSTUS v.3.4.0 (Stanke & Waack, 2003), was trained using the gene structures identified from the transcript-to-genome alignments by TransDecoder v.5.6.0 (Haas et al., 2013). The second predictor, BRAKER v.2.1.6 (Hoff et al., 2019), was trained with the alignments of RNA reads to genomes obtained with Hisat2. Finally, GALBA v.1.0.1 (Brůna et al., 2023) was trained using the proteomes of the three termite and one cockroach species available on InsectBase.

Both the extrinsic evidence (*i.e.*, protein-to-genome and transcript-to-genome alignments) and the *ab initio* gene predictions were combined with EvidenceModeler v.1.1.1 (Haas et al., 2008). The procedure generated consensus gene structures. Genes were not included in the annotation if they contained in-frame stop codons or were inferred by only one *ab initio* predictor and lacked additional evidence from other annotation methods. The consensus gene structures and the transcripts assembled by StringTie (Pertea et al., 2015) were processed with two iterations of PASA v.2.5.2 (Haas et al., 2008), which curated the gene structures, added untranslated regions, and built models for alternative splicing.

The initial annotation of every genome was performed independently, as described above, without considering other genomes sequenced in this study. As a final step, we improved the annotation of every genome with the proteomes of all other termite and cockroach species sequenced here as extrinsic evidence. To do so, we aligned the proteomes of all termite and cockroach species to each genome sequenced in this study using miniprot v.0.7 (Li, 2023). The gene structures identified with these protein-to-genome alignments were combined with the initial annotation with EvidenceModeler v.1.1.1 (Haas et al., 2008). Gene models were filtered and processed with PASA v.2.5.2 (Haas et al., 2008) as described above.

### Removal of microbial contaminations

Although the genomes were sequenced from head tissues to limit contaminations by gut microbes, microbial DNA, such as that of *Wolbachia* and other intracellular parasites, were likely present in our genome assemblies. In order to ensure our 47 genomes were free from microbial contaminations, the proteome of each genome was searched against the non-redundant (nr) database with DIAMOND v2.1.7.161 (Buchfink et al., 2015) with default settings. Homology hits were processed with MEGAN6 (Huson et al., 2007), and each protein was assigned to a taxon based on the best hits. Genomic contigs were excluded when more than 90% of the genes they contained were assigned to Bacteria, Archaea, or viruses.

### Identification of transposons

The transposons were identified using the sensitive mode of EDTA v2.2.0 (Ou et al., 2019). EDTA built a transposon library for every genome by incorporating transposon identification software, including LTRharvest (Ellinghaus et al., 2008), LTR_FINDER_parallel (Ou & Jiang, 2019), and LTR_retriever (Ou & Jiang, 2018) for long terminal repeats (LTRs); Generic Repeat Finder (Shi & Liang, 2019) and TIR-Learner (Su et al., 2019) for terminal inverted repeats (TIRs) and miniature inverted repeat transposable elements (MITEs); HelitronScanner (Xiong et al., 2014) for Helitrons; AnnoSINE (Li et al., 2022) for short interspersed nuclear elements (SINEs); and RepeatModeler (Smit et al., 2015b) for transposons not considered by other programs. Transposons identified with EDTA were soft masked in the genomes with RepeatMasker (Smit et al., 2015a).

### Identification of CAZymes

We analysed the protein sequences from each genome using the run_dbcan python package v. 4.0.0 implemented in dbCAN3 (Zheng et al., 2023). The command run_dbcan assigns genes to CAZyme families with three search tools: HMMER, DIAMOND, and dbCAN_sub. We used these three search tools to annotate our protein sequences. Other parameters were left at their default values. Only genes assigned to a CAZyme family by two or three search tools were retained, as we considered annotations by only one search tool to be unreliable. Finally, we did not consider the Glycosyl Transferases (GTs), as they catalyse the formation of glycosidic bonds (Coutinho et al., 2003) and are not associated with lignocellulose degradation. For the same reasons, we did not consider the Carbohydrate-binding modules (CBM) associated with GT. All other CAZyme classes were considered.

### Maximum-likelihood phylogenetic tree

We reconstructed a phylogenetic tree of termites using 27,610 of the 50,616 termite-specific ultraconserved element (UCE) described in Hellemans et al., 2022a. UCEs were extracted from genome assemblies with the flanking 200 bp at both 5’ and 3’ ends (∼600 bp loci) using PHYLUCE v1.6.6 (Faircloth, 2016) and LASTZ (Harris, 2007) following the procedure described therein. All UCE loci were aligned with MAFFT (Katoh et al., 2002), as implemented in phyluce_align_seqcap_align. Alignments were internally trimmed using phyluce_align_get_gblocks_trimmed_alignments_from_untrimmed that implements Gblocks (Castresana, 2000; Talavera & Castresana, 2007) with default parameters. Loci absent from one or more genomes were filtered out using phyluce_align_get_only_loci_with_min_taxa. The resulting 27,610 alignments were concatenated into a single supermatrix. A maximum-likelihood phylogenetic tree was reconstructed using IQ-TREE v1.6.12 with 1,000 ultrafast bootstrap replicates to assess branch supports (Hoang et al., 2017; Nguyen et al., 2015). We used a GTR model of nucleotide evolution with a gamma-distributed rate of variation across sites and a proportion of invariable sites (GTR+G+I).

### Inferring Hierarchical Orthologous groups

For each gene, we only retained the longest representative isoform, which we used to classify genes into hierarchical orthogroups with OrthoFinder v.2.5.4 (Emms & Kelly, 2019). The program was run using the species tree inferred from UCEs described above as input. Other parameters were left on default settings.

### Time-calibrated phylogenetic tree

We reconstructed the timescale of termite evolution with the program MCMCTREE implemented in the PAML 4.9g package (Yang, 2007). MCMCTREE uses Bayesian estimations of species divergence time with approximate likelihood calculation, an approach that increases the speed and enables the reconstruction of time-calibrated phylogenies with many genetic loci. We used the rooted maximum likelihood phylogenetic tree reconstructed with UCE markers described above as the input tree topology and 1,409 single-copy orthologous genes (SOGs) as sequences. We used 1,409 SOGs instead of 27,610 UCE loci to reduce computational demand to an acceptable level. SOGs were retrieved by selecting hierarchical orthologous groups containing exactly one gene of each of the 47 species sequenced here. SOG protein sequences were aligned using MAFFT v7.520 (Katoh et al., 2002) under auto mode and converted into codon alignments using PAL2NAL v14 (Suyama et al., 2006). SOG codon alignments were concatenated using FASconCAT-G v1.04 (Kück & Meusemann, 2010) and the third codon positions were removed. The final alignment supermatrix was composed of 1.43 million base pairs.

Node calibrations were implemented as lower and upper soft bounds on fourteen internal nodes based on termite and cockroach fossils (Table S16). Justifications for the use of these fossils as node calibrations have been discussed in depth by Bucek et al. (2019, 2022). The probability of the calibrated node age being outside the defined range was set to 2.5%. The maximal root age of the tree was set to 323.2 million years in the MCMCTREE configuration file, which corresponds to the age of *Mylacris anthracophila*, the first roachoid fossil and stem Dictyoptera (Durden, 1969). Fossil ages were retrieved from the Paleobiology Database (https://paleobiodb.org/ accessed October 2023).

We first calculated the gradient and Hessian of the log-likelihood with MCMCTREE using the option “usedata = 3” and the HKY+Gamma model of nucleotide substitution. The calculated values were used for the MCMC sampling of posterior distribution with the approximate likelihood method and the option “usedata = 2”. We used an independent lognormal clock model. The MCMC chain was run for 20,000,000 iterations and sampled every 100 iterations. The first 2,000,000 iterations were discarded as burn-in. We ran the MCMC chain thrice and used Tracer v1.7.1 (Rambaut et al., 2018) to confirm that each chain reached sufficient effective sample size and converged to closely overlap node age interval estimates.

### Evolutionary models for genomic features

We modelled the evolution of eight genomic features using both single-rate and multi-rate likelihood models (Revell & Harmon, 2022). The analyses were performed using our time-calibrated tree. The eight genomic features were: genome size, GC content, transposon content, gene number, CDS number and total length, and intron number and total length. First, we investigated the genomic features of *Cryptocercus meridianus*. We compared (i) two multi-rate models in which distinct rates were given to *Blatta orientalis* + *C. meridianus* and termites to (ii) two multi-rate models in which different rates were given to *B. orientalis* and *C. meridianus* + termites. We defined the selective regimes to model these rates by using one iteration of the stochastic mapping “make.simmap” function of the R package “phytools” (Revell, 2012), under an equal-rates model of state transitions (“ER” argument). We then used the “OUwie” function of the R package “OUwie” (Beaulieu & O’Meara, 2023) to either fit a multi-rate Brownian motion (BM) (*i.e.*, multi *σ^2^*) or a multi-optima Ornstein–Uhlenbeck (OU) (*i.e.*, multi *θ*) models (“BMS” and “OUM” arguments) by supplying the mapped regimes. The likelihood of the four multi-rate models was assessed using the Akaike Information Criterion (AIC).

In a second set of model comparisons, we compared single-rate models of termites with multi-rate models in which “lower” termites (*i.e.*, all extant termite families except Termitidae) and Termitidae, or “lower” termites + Macrotermitinae + Sphaerotermitinae + Foraminitermitinae and other Termitidae, were given distinct rates. *B. orientalis* and *C. meridianus* were omitted from these analyses as the focus was on termites. We used the “fitContinuous” function of the R package “Geiger” (Harmon et al., 2008) to fit four single-rate models: a Brownian motion (BM), an Early-burst model (EB), a time-dependent Pagel’s delta model (δ), and an Ornstein–Uhlenbeck (OU) process. For multi-rate models, we defined the two selective regimes (*i.e.*, Termitidae and “lower” termites, or “lower” termites + Macrotermitinae + Sphaerotermitinae + Foraminitermitinae and other Termitidae) and modelled their evolution using the same methodology as described above. The likelihood of the eight models was assessed using the Akaike Information Criterion (AIC).

### Phylogenetic Generalised Least Squares

We tested for a correlation between eight genome characteristics: genome size, GC content, transposon content, gene number, CDS number and total length, and intron number and total length. We used our 45 termite genomes and the genome of *Cryptocercus meridianus*. To control for the phylogenetic non-independence among species, we carried out Phylogenetic Generalised Least Square (PGLS) analyses. We first calculated the expected covariance under a Brownian model with our time-calibrated phylogenetic tree using the “corBrownian” function implemented in R package ape (Paradis et al., 2004). We then fitted generalised least squares with the correlation structure obtained with “corBrownian” using the “gls” function implemented in nlme (Pinheiro et al., 2024). In total, we performed 28 PGLS analyses and corrected the p-values with FDR calculated with the Benjamini-Hochberg method.

### CAZyme family expansion and contraction

We reconstructed the expansion and contraction of CAZyme families on our termite time-calibrated phylogenetic tree using CAFE5 (Mendes et al., 2020). We first investigated the optimal number of gamma rate categories *K*, which represents the rate of evolution of gene families, by fixing the number of lambda clusters to one (*i.e.*, a single average rate of gene gain and loss for all branches of the tree) and changing the number of gamma rate categories *K* from 1 to 10. Each model was run 50 times to ensure the convergence of lambda. We compared AIC values from the models for which lambda converged to a single estimated value. Models with up to four gamma rate categories *K* yielded convergent values of lambda, with *K* = 4 having the smallest AIC value (AICk=1, L=1 = 4653, AICk=2, L=1 = 4263, AICk=3, L=1 = 4267, AICk=4, L=1 = 4241). We then estimated lambda values for each branch of the tree for *K* = 1. The analyses were repeated between 150 and 250 times for each 92 branches of our phylogenetic tree by allowing one branch at a time to have a unique lambda value differing from that of all other branches of the tree. Repeating the analyses allowed to ensure the convergence of the branch-specific lambda values, which were grouped into two to eight clusters using k-means clustering with the R packages “cluster” (Maechler et al., 2021) and “factoextra” (Kassambara & Mundt, 2017). As a last step, we compared all possible models with one to four gamma rates and one to eight lambda clusters. Each model was run 50 times to test for the convergence of lambda values. The best-fit model, with the lowest AIC value, was the model with two gamma rate categories and two lambda clusters (AICk=2, L=2 = 4198). We re-estimated the branch-specific lambda clustering with *K* = 2, checked for convergence, and found a lower AIC value (AICk=2, L=2 = 4119,04). The selected model and their associated lambda estimates were used to analyse the expansion and contraction of gene families.

### Comparison of wood- and soil-feeding termites using Phylogenetic ANOVA

Our sampling included species feeding on wood and species feeding on soil. Termite diet, either wood or soil, was assigned to each species sequenced in this study based on literature information (Bourguignon et al., 2011; Donovan et al., 2001; Hellemans et al., 2022b) (Table S1). We tested whether diet affects the genomic features and the CAZyme repertoire of termites by comparing wood- and soil-feeding species using phylogenetic ANOVAs. The analyses were performed using our time-calibrated tree. We used the function “phylANOVA” of the R package “phytools” (Revell, 2012) with 10,000 iterations to perform phylogenetic ANOVAs. We performed 53 phylogenetic ANOVA, including eight analyses on genomic features (genome size, GC content, transposon content, gene number, CDS number and total length, and intron number and total length), one analysis on the total number of CAZyme genes found in each genome, and 44 analyses on the abundance (or gene copy number) of 44 CAZyme families. The p-values of the 44 analyses performed on CAZyme families were corrected with the FDR calculated with the Benjamini-Hochberg method. All analyses were repeated twice, once comparing wood-feeding termites + *Cryptocercus meridianus* to soil-feeding termites and once comparing wood-feeding Termitidae to soil-feeding termites, all of which belong to Termitidae.

## Acknowledgements

We thank Kensei Kikuchi, Nobuaki Mizumoto, Juan José González Plaza, Petr Stiblík, Yanli Che, Taisuke Kanao, and Yves Roisin for providing samples and Yukihiro Kinjo for discussion. We thank OIST’s DNA Sequencing Section (SQC) and the Scientific Computation and Data Analysis Section (SCDA) for assistance with sequencing and providing access to the OIST computing cluster, respectively. NGS data production was also carried out at the DRESDEN-concept Genome Center, supported by the DFG Research Infrastructure Program (Project 407482635) and part of the Next Generation Sequencing Competence Network NGS-CN (Project 423957469). The authors thank the HPC Service of FUB-IT (Bennett et al 2020), Freie Universität Berlin, and HPC Service of PALMA, University of Münster, for computing time. This work was supported by subsidiary funding from OIST to TB, including funding for a workshop held at OIST in early December 2022 and funding by the Deutsche Forschungsgemeinschaft (DFG, German Research Foundation) to DPM (MC 436/5-1 and MC 436/7-1) and MCH (HA 8997/1-1). AB was supported by the Czech Science Foundation (GAČR) grant Junior STAR No. 23-08010M. TA was supported by the Japan Society for the Promotion of Science through a DC2 graduate student fellowship. IH was supported by São Paulo State Foundation (FAPESP) grant #2020/08121-5.

## Author Contributions Statement

JŠ, MCH, DPM, and TB conceptualised the experiments. CA, IH, SHel, DSD, ZW, and JŠ collected the samples. CC, CA, and SHe performed lab experiments and generated data. CL, CA, AM, TA, SHel, YMW, AB, and GT analysed the data. TB wrote the original draft manuscript. CL, CA, AM, SHel, YMW, AB, GT, JŠ, MCH, and DPM revised the manuscript. All authors (CL, CA, AM, TA, SHel, YMW, CC, SHe, ZW, IH, DSD, AB, GT, JŠ, MCH, DPM, and TB) read and accepted the final version of this manuscript.

## Compliance with Ethical Standards

Sampling was performed following local and worldwide regulations at the time of collection. Hereafter is a list of permits obtained to sample termite for this project: Brazil, ICMBio/IBAMA (No. 76538/1-3 and No. 75489/1-2) and SISGEN (No. AAE999E, No. A922724 and No. ADA8E5C); Cameroon, No. 000000010/MINRESI/B00/C00/C10/C12 and No. 00000075/MINRESI/B00/C00/C10/C12; French Guiana, No. TREL1902817S/136; Panama, No. SEX/A-36-10. Other samples were collected in regions that did not require permits at the time of sampling.

---

**Table S1.** List of species analysed in this study, with details of family, subfamily, genus, diet, sample code, collection date and location, GPS coordinates, and collectors.

**Table S2.** PromethION long reads information and statistics.

**Table S3.** Statistics of the draft assembly generated with FLYE v2.9 using PromethION long reads.

**Table S4.** Statistics of genome assemblies polished with short reads using HyPo v1.0.3. and collapsed into a single haplotype using purge_haplotigs v1.1.2.

**Table S5.** List of transcriptomes used to annotate the genome assemblies.

**Table S6.** Statistics of genome assemblies scaffolded with Hi-C and OMNI-C data.

**Table S7.** List of genome characteristics analysed in this study.

**Table S8.** Results of four models representing the evolution of genome characteristics in termites and two cockroaches. Two models used two rates of evolution for cockroaches and termites and two models used two rates of evolution for *Blatta* and *Cryptocercus* + termites.

**Table S9.** Results of eight models representing the evolution of genome characteristics in termites. Four models used two rates of evolution for “lower” termites and Termitidae or “lower” termites + Macrotermitinae + Sphaerotermitinae + Foraminitermitinae and other Termitidae and four models used a single rate of evolution for all termites.

**Table S10.** Results of the Phylogenetic Generalised Least Squares analyses performed on all possible pairwise comparisons of eight genomic features.

**Table S11.** Results of the Phylogenetic Generalised Least Squares analyses performed on genome assembly size and four measures of gene number.

**Table S12.** Results of the Phylogenetic ANOVAs comparing *Cryptocercus* + wood-feeding termites to soil-feeding termites and wood-feeding Termitidae to soil-feeding termites for eight genomic features.

**Table S13.** List of CAZyme families and their copy number in all genomes sequenced in this study.

**Table S14.** Results of the analysis of CAZyme family gene expansion and contraction with CAFE5.

**Table S15.** Results of the Phylogenetic ANOVAs comparing *Cryptocercus* + wood-feeding termites to soil-feeding termites and wood-feeding Termitidae to soil-feeding termites for the total number of CAZyme genes found in each genome and for the gene copy number of 44 CAZyme families.

**Table S16.** List of fossils used as calibration for estimating divergence time.

